# Interface-Resolved Proteomics of Cell–Cell Membranes Reveals Early Spatial Polarity in a Vertebrate Embryo

**DOI:** 10.64898/2026.01.10.698819

**Authors:** Fei Zhou, Peter Nemes

## Abstract

Cell–cell membrane interfaces are central sites of adhesion, signaling, and polarity establishment, yet they have remained inaccessible to proteome-wide analysis as discrete analytical units. Here, we report an interface-resolved proteomics workflow that isolates intact intercellular membrane segments from single, identified blastomeres and quantitatively profiles their protein composition. Using a microdissection-enabled strategy combined with optimized mild-detergent extraction and high-sensitivity high-resolution mass spectrometry, we achieve deep coverage of low-input membrane samples, identifying ∼3,000 proteins per interface type, including over 100 annotated plasma-membrane proteins. Applying this approach to defined dorsal–dorsal, dorsal–ventral, and ventral–ventral cell–cell interfaces in a 16-cell chordate embryo model, *Xenopus laevis*, reveals reproducible, interface-specific proteomic signatures that distinguish neighboring membrane contacts along the primary body axis. Region-enriched proteins include regulators of membrane trafficking, signaling, cytoskeletal organization, and metabolic pathways linked to early dorsal–ventral patterning. These results demonstrate that intercellular membrane interfaces exhibit molecular polarity at early developmental stages and establish interface-resolved proteomics as a general strategy for mapping spatially organized biochemical activities at cell–cell contacts.

## INTRODUCTION

Cell–cell membrane interfaces are fundamental sites where adhesion, signaling, and polarity cues are integrated to coordinate cellular behavior. These nanoscale regions host adhesion complexes, receptors, transporters, and cytoskeletal linkers that collectively organize communication between neighboring cells. During early embryonic development, such interfaces play a critical role in transmitting positional information that underlies axis formation and cell fate decisions. In amphibian embryos, dorsal–ventral membrane contacts have been implicated in noncanonical signaling pathways that contribute to early body plan patterning.^1–4^ Despite their biological importance, the molecular composition of intercellular membrane interfaces has remained largely unexplored.

Proteomic analysis of cell membranes has advanced substantially over the past two decades. Conventional surface and subcellular proteomics workflows enrich whole plasma membranes or broad membrane fractions from large numbers of cells and have provided valuable insight into global membrane organization and dynamics.^5–9^ However, these approaches, often grounded in mass spectrometry (MS), do not resolve specific membrane regions formed between identified neighboring cells and therefore cannot interrogate the molecular heterogeneity that may exist between distinct cell–cell contacts within the same tissue or embryo. For intercellular interfaces, this limitation is compounded by their small physical footprint, low protein content, and close association with cytoplasmic and yolk-rich material. As a result, cell–cell membrane interfaces have not been accessible as discrete analytical units for high-resolution MS (HRMS) based proteomics.^5–6^

Recent advances in subcellular proteomics have improved sensitivity and coverage for limited-input samples through optimized extraction chemistries, streamlined workflows, and enhanced MS detection.^10–13^ Mild nonionic detergents, low-loss sample handling, and improved data acquisition strategies now support quantitative analysis of nanogram-scale proteomes, including membrane-associated proteins.^14–19^ These developments have enabled deep proteomic profiling of eggs, embryos, and isolated membrane microdomains, revealing extensive remodeling of signaling and trafficking pathways during early development.^20–23^ In parallel, progress in microsurgical sampling and *in situ*/vivo microextraction has extended proteomics and metabolomics to defined cellular and subcellular regions.^3–4, 15, 20–21, 24–26^ Nevertheless, isolating intact intercellular membrane segments from living embryos remains technically challenging, particularly as cells adhere tightly, decrease in size, and divide rapidly during development.

The vertebrate embryo *Xenopus laevis* provides a uniquely tractable system for addressing this analytical challenge. Its large, stereotypically arranged blastomeres can be reliably identified using established fate maps and are sufficiently robust for direct microsurgical manipulation.^12, 27–29^ These features have made *Xenopus* an enabling platform for the development and validation of single-cell and subcellular MS workflows.^12, 30^ At the same time, the early embryo presents stringent analytical constraints, including high yolk content, limited material per interface, and complex three-dimensional geometry. These characteristics provide a demanding but informative testbed for developing strategies that render intercellular membrane interfaces accessible to quantitative proteomic analysis.

Here, we report an interface-resolved proteomics workflow that isolates and quantitatively profiles intact intercellular membrane segments between single, identified blastomeres. Using a microdissection-enabled strategy combined with optimized mild-detergent extraction and high-sensitivity HRMS, we convert cell–cell membrane interfaces into reproducible analytical units. We apply this approach to dorsal–dorsal (DD), dorsal–ventral (DV), and ventral–ventral (VV) cell–cell interfaces at the 16-cell stage of a vertebrate embryo to assess whether distinct membrane contacts exhibit measurable proteomic differences. The resulting interface-resolved proteome maps reveal reproducible spatial heterogeneity consistent with early membrane polarity and establish a general framework for probing the molecular organization of cell–cell contacts in developing systems.

## RESULTS AND DISCUSSION

### Interface-Resolved Isolation of Intercellular Membrane Segments

Our experimental strategy couples precision microdissection of early embryos with high-sensitivity bottom-up nano-flow liquid chromatography (nanoLC) HRMS to profile defined intercellular membrane interfaces (**Fig. 1**). In the early vertebrate embryo, cell–cell contacts are thin, curved membrane sheets embedded within yolk-rich cytoplasm, making them difficult to excise intact and prone to mechanical damage. Direct isolation of these interfaces, therefore, requires careful control of embryo geometry and tissue handling. We focused on isolating membrane segments from three defined cell–cell interfaces at the 16-cell stage: D1.1–D1.2 (D11–D12, or DD), D1.2–V1.2 (D12–V12, or DV), and V1.2–V1.1 (V12–V11, or VV). These interfaces demarcate spatial communication pathways along the primary D–V body axis and provide a controlled testbed for assessing whether neighboring membrane contacts differ in molecular composition. Following removal of the vitelline membrane with a pair of fine forceps, embryos were oriented to expose the animal hemisphere and gently flattened to make intercellular junctions accessible for microsurgery (**Fig. 2**).

**Figure 1.**
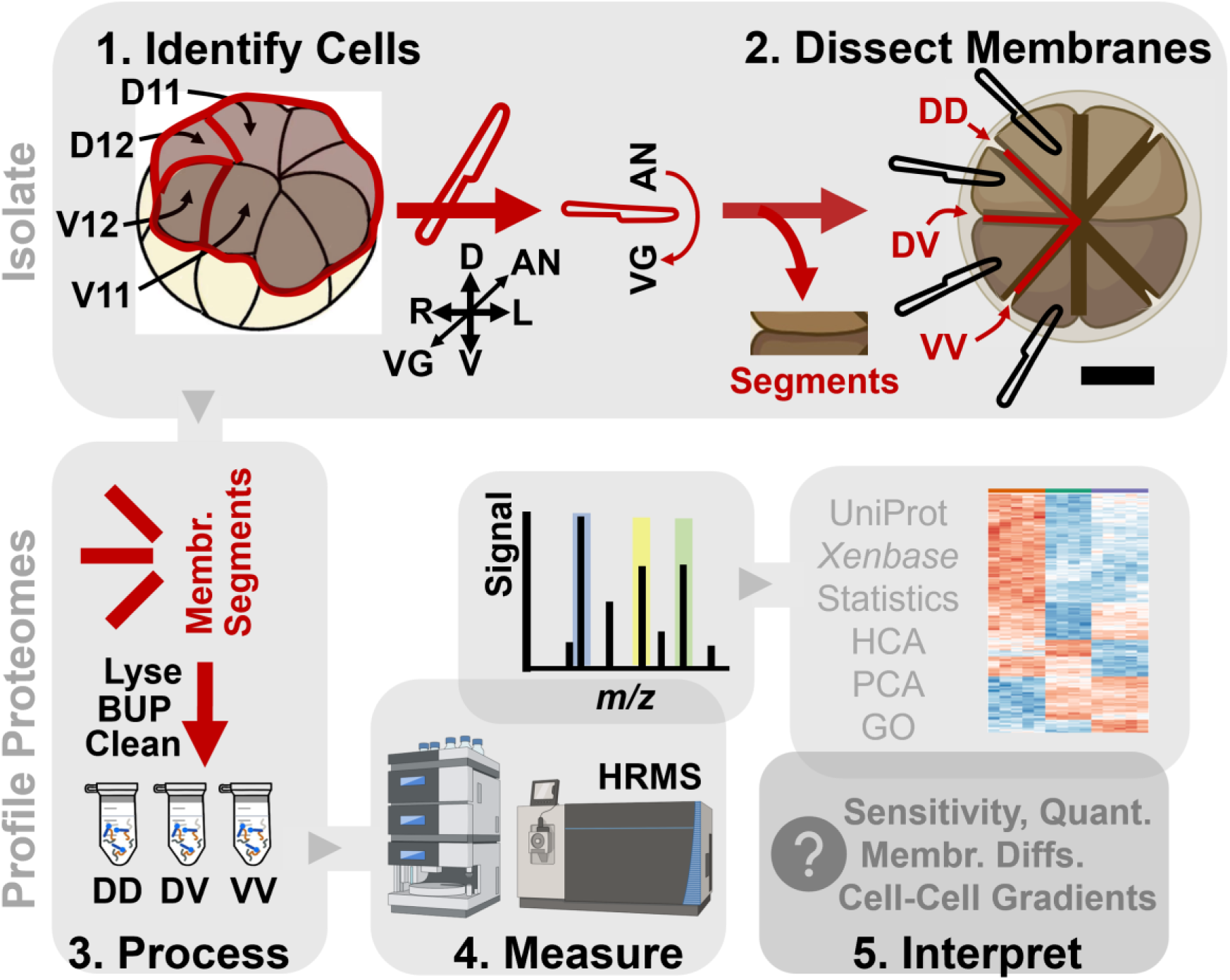
Workflow steps for interface-resolved membrane proteomics in *X. laevis* embryos. *(1)* Identify cells. D11, D12, V12, and V11 blastomeres are identified in the 16-cell embryo along the animal–vegetal (AN–VG), dorsal–ventral (D–V), and left–right (L–R) axes. *(2)* Dissect interfaces. Precision microdissection exposes and excises ∼300-µm segments from the D11–D12 (DD), D12–V12 (DV), and V12–V11 (VV) intercellular membrane interfaces with minimal cytoplasmic carryover. *(3)* Process. Interface segments are pooled by interface type and prepared for bottom-up proteomics (BUP) using an optimized mild-detergent workflow compatible with low-input membrane samples. *(4)* Measure. Peptides are analyzed by reversed-phase nanoLC and HRMS. *(5)* Interpret. Quantitative DD, DV, and VV interface proteomes are evaluated using multivariate and statistical analyses, including principal component analysis (PCA) and hierarchical cluster analysis (HCA), and annotated using UniProt, Xenbase, Gene Ontology (GO), and STRING to support hypothesis generation about spatially organized signaling and metabolism. (Partially created with BioRender.)

**Figure 2.**
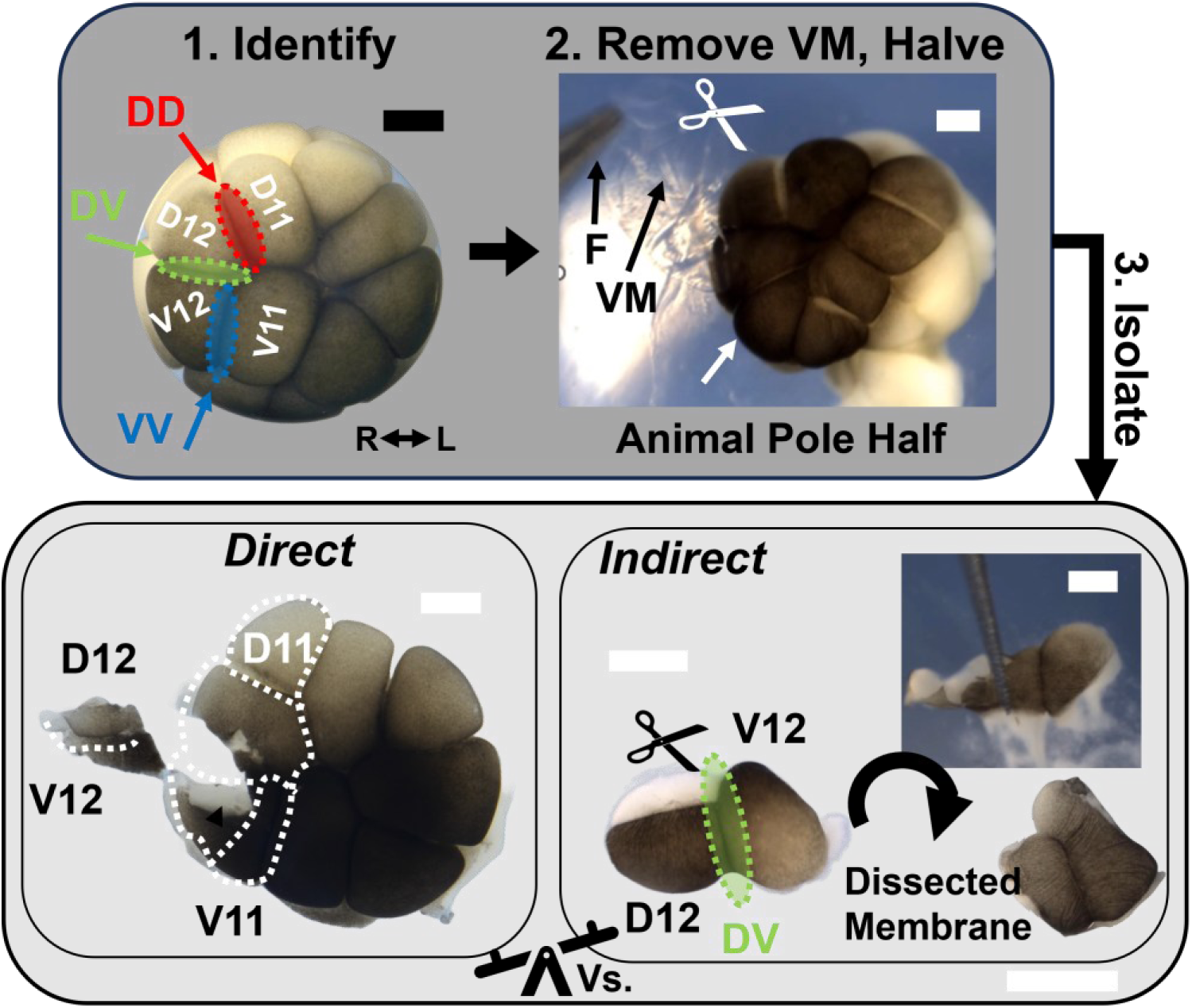
Microdissection of intercellular membrane interfaces in *X. laevis* embryos. Removal of the vitelline membrane (VM) from the vegetal side using sharpened forceps (F) exposed the D11–D12 (DD), D12–V12 (DV), and V12–V11 (VV) cell–cell interfaces in the 16-cell embryo for microsurgery. Direct approach: DD, DV, and VV interfaces were sectioned *in situ* from the half embryo, yielding ∼50% intact segments due to tearing and adhesion. Indirect approach: neighboring cells were dissected away, their contents were gently peeled back, and ca. three-fourths of the exposed membrane sheet was excised as a ∼300 µm segment, increasing success to ∼70%. Representative isolated membrane segments are shown. Scale bars, 250 µm.

We initially evaluated a “direct” excision approach in which the target interface was sectioned *in situ* from the half embryo by inserting a microscalpel between adjacent blastomeres. Although this method preserved anatomical context, it yielded intact membrane segments in only approximately half of the attempts (5 out of 10 trials), with frequent failures arising from tearing or adhesion of the delicate membrane to surgical tools. To mitigate adhesion and improve handling, we supplemented the dissection setup with small droplets of glycerol on the agar surface and into the dissection medium, which promoted membrane stability and reduced sticking during excision and transfer. However, tissue disruption and membrane damage remained frequent and the overall recovery did not improve sufficiently, limiting the utility of the direct method.

To overcome these limitations, we developed an “indirect” isolation strategy in which the cells flanking the target interface were first dissected away, leaving the intercellular membrane sheet exposed and mechanically supported (**Fig. 2**). The contents of neighboring blastomeres were gently peeled back, allowing the interface to be excised with minimal stress. This approach increased the recovery of intact membrane segments to ∼70% and enabled routine isolation of ∼300-µm-long interface segments suitable for downstream proteomic analysis. The improved yield and reproducibility of the indirect strategy allowed scalable collection of DD, DV, and VV membrane interfaces from multiple embryos for quantitative profiling.

### Optimization of Protein Extraction and Solubilization for Low-Input Membrane Interfaces

Sensitive proteomic profiling of microdissected intercellular membrane interfaces required efficient solubilization of membrane-associated proteins while minimizing sample loss from limited input material. We therefore systematically evaluated extraction strategies commonly used in membrane proteomics to determine their suitability for sub-microgram samples enriched in yolk-derived background (**Table S1**).

As a benchmark, we first assessed a classical ultracentrifugation (UF)-based membrane enrichment workflow using pooled embryos (**Fig. 3**). When applied to large starting material, this approach yielded deep proteome coverage, identifying ∼4,350 proteins, including nearly 500 gene ontology (GO)-annotated membrane proteins. However, this performance did not scale to the limited inputs obtainable from microdissected intercellular interfaces. While membrane pellets were readily recovered from preparations starting with 50–10 embryos, scaling to half an embryo or smaller inputs produced pellets that were difficult to isolate cleanly and were enriched to ∼50% yolk content, identifying fewer than ∼85 membrane proteins. These limitations likely arise from both physical handling constraints and nonspecific losses during multiple transfer steps, rendering UF poorly suited for interface-level proteomics in our study.

**Figure 3.**
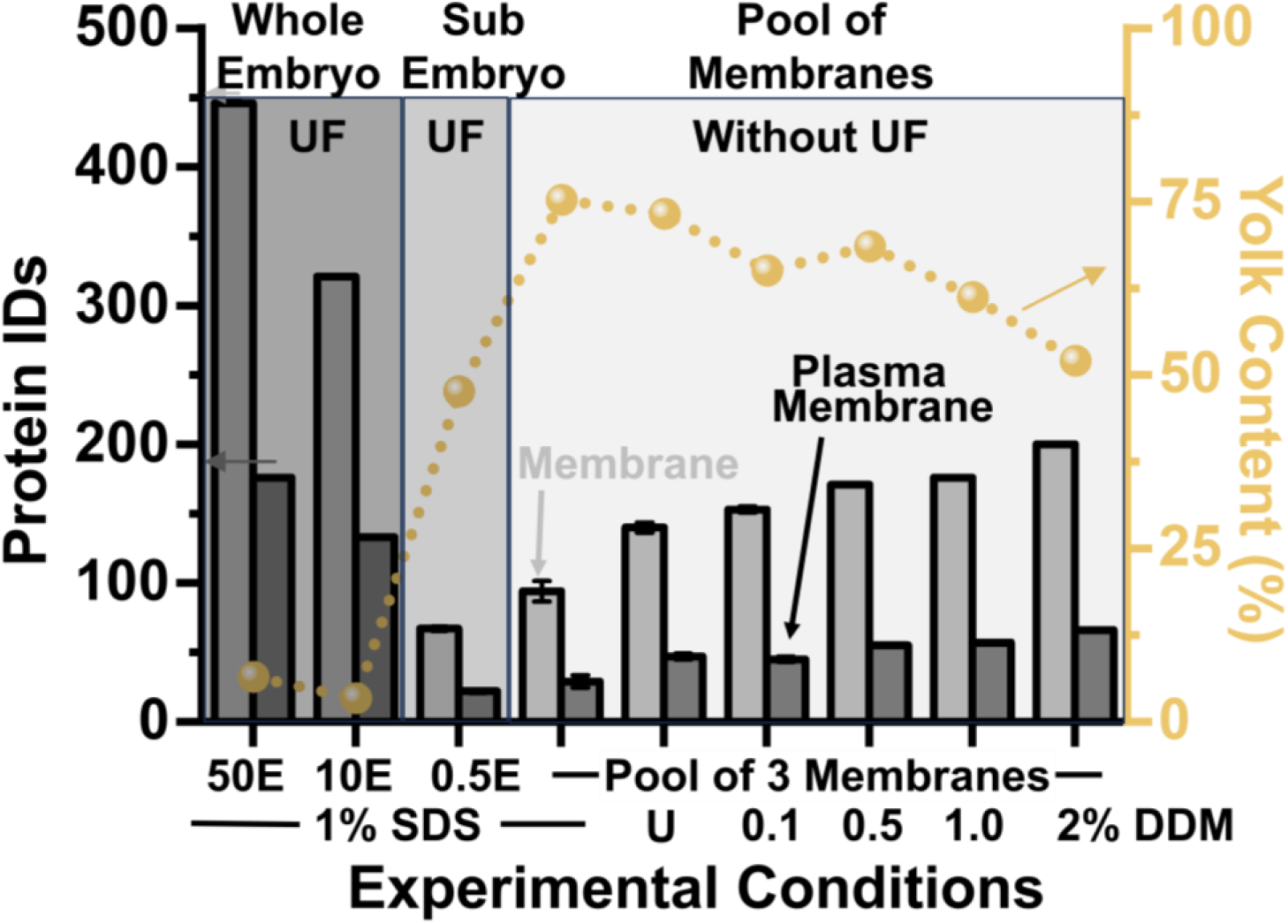
Optimization of membrane protein extraction and solubilization. Comparison of classical ultracentrifugation (UF) vs limited-input extractions using 8 M urea (U), SDS, or 0.1–2% DDM. Vitelline membrane-removed embryos (Es) were pooled to benchmark UF performance, whereas pools of three ∼300-µm membrane segments were used to test limited-input conditions. Each preparation yielded ∼20 µg digest and was analyzed in technical triplicate. Identified proteins are listed in **Table S1**, with Gene Ontology-annotated membrane proteins curated in **Table S2**. The comparison summarizes recovery of membrane proteins (MP), plasma-membrane proteins (PMP), and the yolk-associated proteins (notably Vtga2, Vtga2.L, Vtgb1.L, Vtgb1.S). For low-input samples, 2% DDM without UF provided the deepest coverage of membrane proteins despite a higher yolk background. Error bars, standard deviation (3 technical replicates), within symbols.

We therefore evaluated UF-free extraction strategies that minimize sample handling. Chaotropic denaturation with urea (8 M) was compared to solubilization with a strong ionic (SDS) and mild nonionic (n-dodecyl-β-D-maltoside, DDM) detergent using pooled intercellular membrane segments (**Fig. 3**). Because yolk proteins constitute the majority, ∼90%, of the early embryonic proteome, posing a dominant background interference for MS detection,^31–33^ effective separation of membrane proteins from yolk-derived background was a critical consideration. Urea and SDS provided strong solubilization but required additional cleanup steps before nanoLC–HRMS analysis, which introduced losses at low input.

Among the conditions tested, solubilization with 2% DDM without UF provided the most effective balance between membrane protein recovery and workflow simplicity. To maintain compatibility with downstream chromatography, residual detergent was mitigated by incorporating a brief high-organic wash at the end of each nanoLC gradient.^34^ This approach enabled identification of ∼230 membrane proteins from pools of 3 intercellular membrane segments, including 86 annotated plasma-membrane proteins (**Fig. 3B**; **Table S2**). These results demonstrate that mild-detergent extraction without UF is essential for preserving membrane proteome coverage from limited intercellular interface material.

### Selection of HRMS Acquisition Strategy for Low-Input Membrane Proteomes

To maximize proteome coverage from limited intercellular membrane material, we next evaluated data acquisition strategies suitable for low-input samples. Both data-dependent acquisition (DDA) and data-independent acquisition (DIA) have been applied successfully in subcellular and single-cell proteomics, but their relative performance depends strongly on sample complexity, input amount, and chromatographic separation time. A pooled digest generated from 20 DV membrane segments was used to construct a spectral library through high-pH fractionation followed by DDA analysis of each. Using this library containing >5,000 proteins, we compared library-based and library-free DIA workflows across multiple quadrupole-isolation window schemes for sequencing (10, 15, or 25 Th widths) while maintaining a fixed acquisition cycle time (3 s). Across all DIA configurations tested, ∼1,400 proteins were identified from 500 ng of digest (**Fig. 4A**), including 89 membrane proteins and 29 annotated plasma-membrane proteins (**Fig. 4B**).

**Figure 4.**
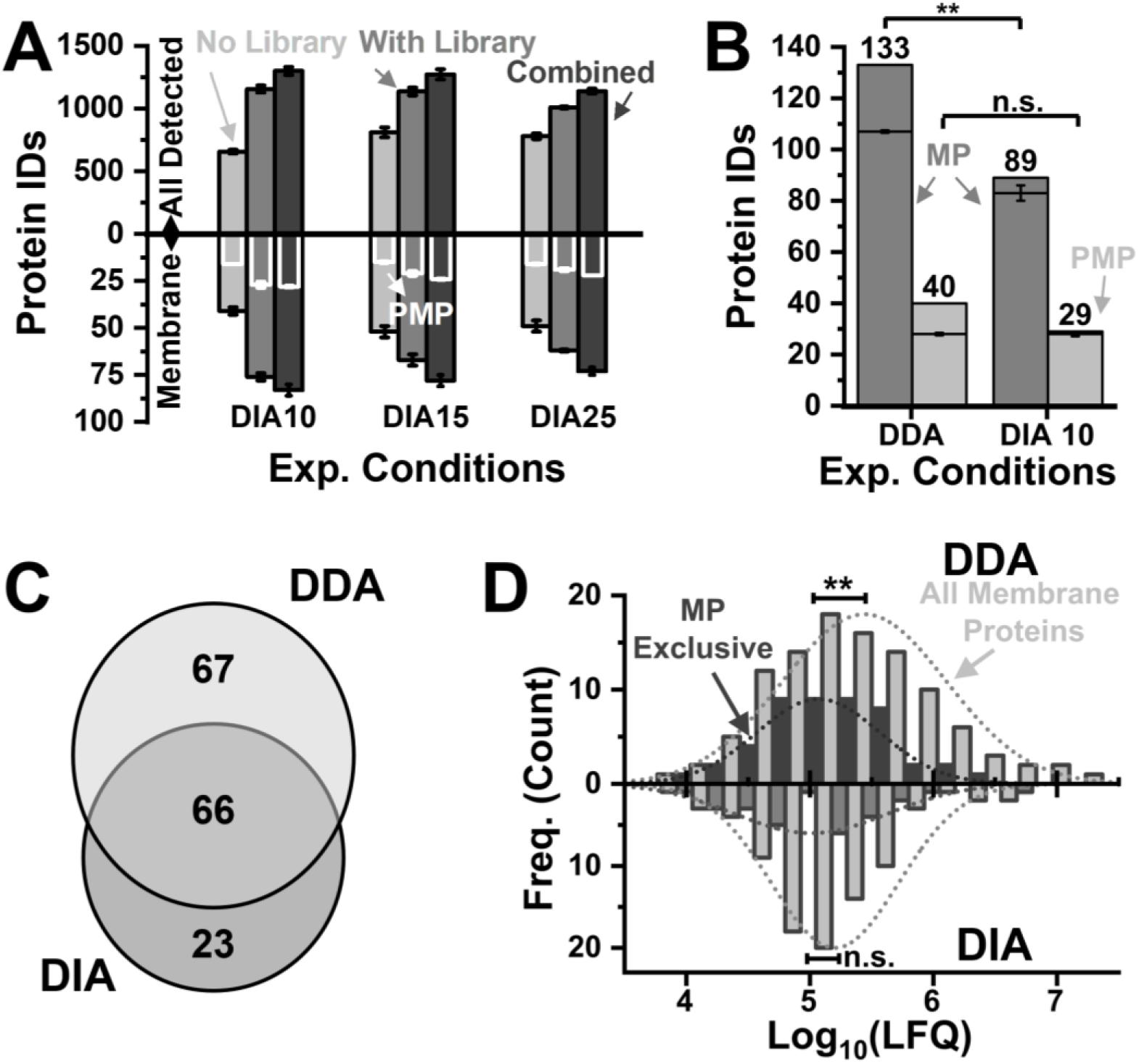
Tailoring HRMS acquisition for low-input membrane proteomics. A pooled digest from n = 20 DV membrane segments was analyzed using data-dependent acquisition (DDA) and data-independent acquisition (DIA). **(A)** DIA performance across quadrupole isolation windows (25, 15, 10 Th) and processing modes (library-based, library-free, combined). DIA with 10 Th provided the greatest depth under these conditions. White bars denote plasma-membrane proteins (PMP). **(B)** Comparison of identified membrane proteins (MP) and PMPs for DIA and DDA and the corresponding **(C)** Venn diagrams for overlap. **(D)** LFQ protein intensity distributions for membrane proteins detected uniquely by each method. DDA captured additional low-abundance MPs. Statistics: Error bars, standard deviation (3 technical replicates); **, *p* < 0.01, Mann–Whitney U test; n.s., not significant.

Under identical sample-loading and separation conditions, DDA provided deeper coverage of membrane-associated proteins, identifying 133 membrane proteins in cumulative analyses (**Fig. 4B,C**; **Table S3**). Notably, membrane proteins detected exclusively by DDA populated the lower-abundance range of the label-free quantification (LFQ) distributions approximating protein concentration (**Fig. 4D**), indicating improved sensitivity for low-abundance species. This advantage through DDA likely reflects the long chromatographic separation window used here (∼4 h/run), which enabled extensive precursor peptide sequencing and favored detection of scarce membrane-derived peptides. Based on these results, we selected DDA for quantitative profiling of intercellular membrane interfaces in subsequent experiments. While DIA offers advantages for throughput and reproducibility in higher-input regimes, DDA provided superior sensitivity for deep characterization of low-input membrane proteomes under the conditions required for interface-resolved analysis.

### Spatial Proteomic Profiling of Intercellular Membrane Interfaces

We next applied the interface-resolved proteomics workflow to investigate whether distinct intercellular membrane contacts exhibit reproducible molecular differences during early embryonic development. We focused on DD, DV, VV membrane interfaces at the 16-cell stage (**Fig. 5A**), which demarcate spatial communication pathways along the primary D–V axis.^35–36^ For each interface type, ∼3 membrane segments of ∼300 µm length were pooled from different embryos and processed as biological replicates. Approximately 500 ng total proteome digest, containing <250 ng of yolk-free proteome (**Fig. 3**) per interface type was analyzed by nanoLC–HRMS in technical triplicate.

**Figure 5.**
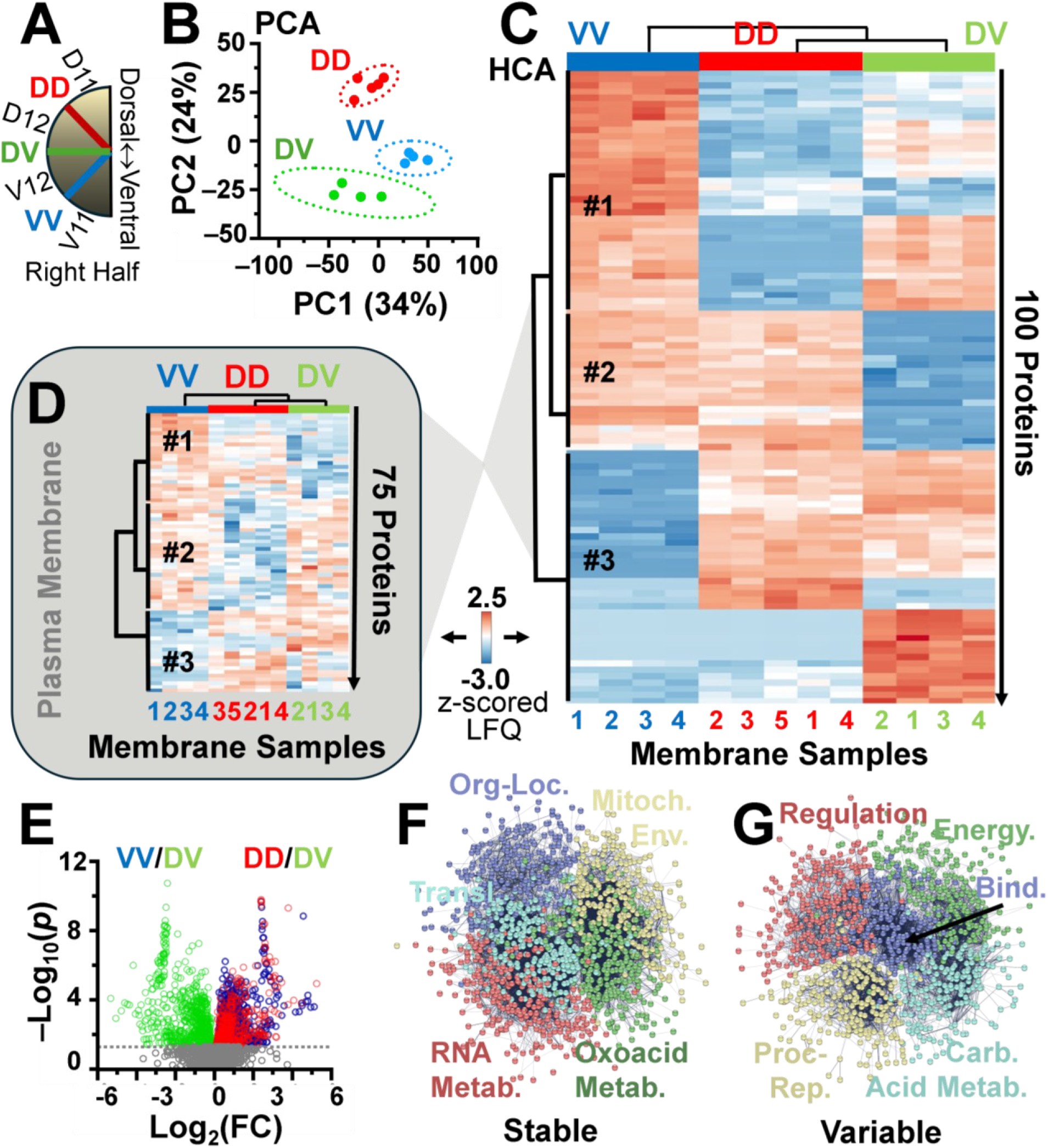
Spatial intercellular membrane proteomes in 16-cell *X. laevis* embryos. **(A)** Orientation schematic showing the dorsal–ventral arrangement of the sampled intercellular interfaces on the right embryonic hemisphere. DD denotes D11–D12, DV denotes D12–V12, and VV denotes V12–V11 cell–cell interfaces. **(B)** Unsupervised PCA of the LFQ protein intensities separates samples into three clusters corresponding to VV, DD, and DV interfaces (dashed ellipses, 95% confidence regions; PC1 and PC2 explained variance indicated on axes). **(C)** HCA–heatmap of the top 100 most variable proteins across interfaces (rows, z-scored LFQ; columns, membrane samples), showing interface-specific abundance patterns and clustering by interface identity. **(D)** HCA heatmap of the 75 most variable plasma-membrane proteins (rows, z-scored LFQ; columns, membrane samples), demonstrating that spatial separation is preserved within the plasma-membrane subset (color scale shown). **(E)** Volcano plot summarizing differential abundance for VV vs DV and DD vs DV comparisons (independent axis, log2 fold change; dependent axis, −log10(*p*); dashed line indicates *p* = 0.05, two-sided Student’s t-test). **(F)** STRING protein–protein interaction network for proteins classified as stable across interfaces (*p* ≥ 0.05), with functional group labels indicating major enriched modules. **(G)** STRING network for proteins classified as variable across interfaces (*p* < 0.05), highlighting interface-associated modules. Key: Bind., binding; Metab., metabolism; Mitoch. Env., mitochondrial envelope; Org-Loc., organization–localization; Proc.-Rep., processing–repair.

Across the 3 interface types, up to ∼3,000 proteins were identified, of which ∼2,600 were quantified (**Table S4**). To our knowledge, this dataset represents the first proteome-wide characterization of intact intercellular membrane segments in a vertebrate embryo. Unsupervised principal component analysis (PCA) of log-transformed, normalized LFQ intensities separated the samples (n = 13 biological replicates) into 3 well-defined clusters along the first two principal components accounting for ∼57% of total variance in the data (**Fig. 5B**). Upon revealing the identities, these sample groups corresponded to DD, DV, and VV interfaces. These results indicate that membrane proteome composition differs systematically between the neighboring cell–cell contacts.

Unsupervised hierarchical cluster analysis (HCA) of the 100 most variable proteins across all samples revealed a similar organization, with biological replicates clustering by interface identity rather than by embryo of origin (**Fig. 5C**; **Fig. S1**). Importantly, restricting the analysis to GO-annotated plasma-membrane proteins preserved this spatial separation. HCA of the 75 most variable plasma-membrane proteins clearly distinguished DD, DV, and VV interfaces (**Fig. 5D**; **Fig. S2**), demonstrating that molecular heterogeneity is maintained at the level of membrane-embedded proteins rather than being driven solely by cytosolic carryover from dissections. Combined, these results show that intercellular membrane interfaces exhibit reproducible, interface-specific proteomic signatures at an early developmental stage.

### Functional Organization of Interface-Specific Membrane Proteomes

To further interpret the molecular differences among intercellular membrane interfaces, we performed pairwise statistical comparisons of protein abundances between DD, DV, and VV membranes. Differential abundance analysis identified both shared and interface-enriched proteins, revealing systematic variation in membrane composition across the D–V axis (**Fig. 5E**). These comparisons indicate that while a substantial fraction of the interface proteome is conserved, distinct subsets of proteins are preferentially associated with specific membrane contacts.

To distinguish broadly shared functions from spatially variable ones, proteins were classified based on statistical significance as stable (*p* ≥ 0.05) or variable (*p* < 0.05) in abundance across interfaces. Known and predicted protein–protein associations were then queried using STRING to assess whether these groups organized into coherent functional modules (**SI Methods**). Proteins with stable abundance across DD, DV, and VV interfaces formed densely connected networks enriched for processes related to RNA metabolism, mitochondrial inner-membrane organization, protein translation, and general cellular organization–localization functions (**Fig. 5F**). GO and KEGG pathway annotations linked these proteins to cytoskeletal organization, endocytosis, tight junctions, and protein processing pathways, consistent with housekeeping and shared structural roles at cell–cell contacts.

In contrast, proteins exhibiting interface-specific abundance differences organized into networks associated with more specialized molecular functions (**Fig. 5G**). One prominent cluster, annotated as carboxylic acid metabolism, was enriched for pathways involved in carbon metabolism, valine–leucine–isoleucine degradation, pyruvate metabolism, glycolysis and gluconeogenesis, the tricarboxylic acid cycle, glycine–serine–threonine metabolism, and oxidative phosphorylation. Additional clusters mapped to protein processing and repair, including the proteasome, protein processing in the endoplasmic reticulum, DNA replication, mismatch repair, and ubiquitin-mediated proteolysis, as well as to regulatory pathways such as endocytosis, phagosome formation, tight junctions, and gap junctions. A binding-associated cluster was enriched for ribosomal components, RNA degradation, amino acid biosynthesis, and glycine–serine–threonine metabolism, whereas an energy-related cluster contained numerous components of the electron transport chain and pathways linked to oxidative phosphorylation, the pentose phosphate pathway, and glutathione metabolism.

Notably, several of the interface-enriched metabolic pathways converge on glycine–serine–threonine metabolism. Our previous single-cell metabolomics studies identified metabolites within this pathway that exhibit moonlighting activities and influence dorsal–ventral patterning in the early embryo.^3^ While the present proteomic data do not establish causal relationships, the spatial organization of enzymes and regulatory proteins linked to these metabolic pathways at defined cell–cell interfaces suggests that intercellular membrane contacts may serve as localized platforms for coordinating metabolic and signaling activities, potentially acting in “moonlighting”, or noncanonical roles to pattern the vertebrate embryonic body plan. These findings provide a biochemical framework for generating testable hypotheses that connect interface-resolved proteome polarity with metabolite-driven regulation of early embryonic patterning.

## CONCLUSIONS

We have established an interface-resolved proteomics workflow that enables quantitative analysis of intact intercellular membrane segments between single, identified blastomeres in a developing vertebrate embryo. By integrating precision microdissection with mild-detergent extraction and high-sensitivity nanoLC–HRMS, this approach converts cell–cell membrane interfaces into reproducible analytical units that are compatible with deep proteomic profiling. Using this strategy, we achieved identification of ∼3,000 proteins per interface type, including extensive coverage of annotated plasma-membrane proteins, from limited interfacial material. To our knowledge, this work provides the first proteome-wide characterization of cell–cell membrane segments in a chordate embryo and the deepest membrane-annotated proteome reported to date for early *X. laevis* development.

Application of this workflow to DD, DV, and VV membrane interfaces at the 16-cell stage revealed reproducible and interface-specific proteomic signatures. These differences indicate that molecular polarity is already present at intercellular membrane contacts early in embryonic development. Interface-enriched proteins mapped to functional modules involved in membrane trafficking, signaling, cytoskeletal organization, and metabolism, including pathways associated with glycine–serine–threonine metabolism that have previously been implicated in dorsal–ventral patterning at the single-cell level.

Although the present study does not establish causal roles for individual proteins or pathways, the spatial organization of these molecular features at defined cell–cell interfaces suggests that membrane contacts may act as localized hubs for coordinating biochemical activities during early development. By revealing proteome-scale heterogeneity at intercellular interfaces, this work provides a foundation for testing how interface-specific molecular organization contributes to embryonic pattern formation. These interface-resolved proteomes create new opportunities to investigate the biochemical mechanisms that contribute to early pattern formation and cell communication and provide a resource for designing future functional studies at defined membrane contacts.

More broadly, this study expands the scope of spatial proteomics by rendering intercellular membrane interfaces accessible to quantitative analysis. The workflow is compatible with low-flow separations, post-translational modification–selective enrichments, and multiplexed quantification strategies, and it is adaptable to other developing systems and tissues. Interface-resolved proteomics, therefore, offers a general framework for interrogating the molecular architecture of cell–cell contacts and for linking spatially organized proteomes to developmental and physiological function.

## Disclosures

The authors declare no competing interests.

## AUTHOR CONTRIBUTIONS

P.N. conceptualized the study and designed the objectives. F.Z. isolated, processed, and measured the membrane proteomes. F.Z. and P.N. analyzed the data and prepared the draft report. P.N. edited and finalized the report. P.N. obtained the funding. Both authors approved the final report.

## ACKNOWLEDGMENTS

This research was supported by the National Institute of General Medical Sciences of the National Institutes of Health (award no. 2R35GM1247555 to P.N.). We thank Sergei Sukharev (Department of Biology, University of Maryland, College Park) for providing access to the ultracentrifuge and assisting with its operation.

## TABLE-OF-CONTENT IMAGE

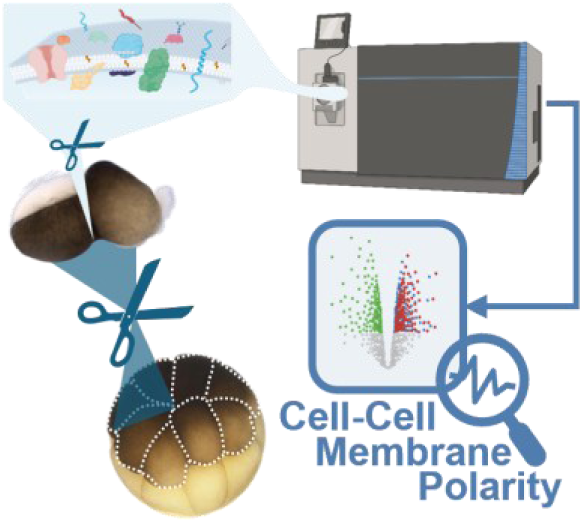

***TOC Image Description*.** A microdissection-enabled HRMS workflow isolates intact intercellular membrane interfaces from single, identified *Xenopus* blastomeres and quantifies their proteomes, revealing interface-specific signatures consistent with early membrane polarity and enabling interface-resolved spatial proteomics.

## Notes

https://www.ebi.ac.uk/pride/

### Competing Interest Statement

The authors have declared no competing interest.

## REFERENCES

1. Shi, D. L., Canonical and non-canonical Wnt signaling generates molecular and cellular asymmetries to establish embryonic axes. J. Dev. Biol. 2024, 12 (3), No. jdb12030020.

2. Goutam, R. S.; Kumar, V.; Lee, U.; Kim, J., Bone morphogenetic protein-mediated Ventx3.2 regulates mesendoderm patterning and gut morphogenesis during *Xenopus* embryogenesis. Biochem. Biophys. Res. Commun. 2025, 784, No. 152623.

3. Onjiko, R. M.; Moody, S. A.; Nemes, P., Single-cell mass spectrometry reveals small molecules that affect cell fates in the 16-cell embryo. Proc. Natl. Acad. Sci. U. S. A. 2015, 112 (21), 6545–6550.

4. Lombard-Banek, C.; Li, J.; Portero, E. P.; Onjiko, R. M.; Singer, C. D.; Plotnick, D. O.; Al Shabeeb, R. Q.; Nemes, P., *In vivo* subcellular mass spectrometry enables proteo-metabolomic single-cell systems biology in a chordate embryo developing to a normally behaving tadpole (*X. laevis*). Angew. Chem. Int. Ed. 2021, 60 (23), 12852–12858.

5. Bhushan, V.; Nita-Lazar, A., Recent advancements in subcellular proteomics: growing impact of organellar protein niches on the understanding of cell biology. J. Proteome Res. 2024, 23 (8), 2700–2722.

6. Labib, M.; Kelley, S. O., Single-cell analysis targeting the proteome. Nature Reviews Chemistry 2020, 4 (3), 143–158.

7. Yang, H.-C.; Li, W.; Sun, J.; Gross, M. L., Advances in mass spectrometry on membrane proteins. Membranes 2023, 13 (5), No. 457.

8. Helbig, A. O.; Heck, A. J. R.; Slijper, M., Exploring the membrane proteome-challenges and analytical strategies. J. Proteom. 2010, 73 (5), 868–878.

9. Wu, C. C.; Yates, J. R., The application of mass spectrometry to membrane proteomics. Nat. Biotechnol. 2003, 21 (3), 262–267.

10. Christopher, J. A.; Stadler, C.; Martin, C. E.; Morgenstern, M.; Pan, Y. B.; Betsinger, C. N.; Rattray, D. G.; Mahdessian, D.; Gingras, A. C.; Warscheid, B.; Lehtiö, J.; Cristea, I. M.; Foster, L. J.; Emili, A.; Lilley, K. S., Subcellular proteomics. Nature Reviews Methods Primers 2021, 1 (1), No. s43586-021-00029-y.

11. Arslan, T.; Pan, Y. B.; Mermelekas, G.; Vesterlund, M.; Orre, L. M.; Lehtiö, J., SubCellBarCode: integrated workflow for robust spatial proteomics by mass spectrometry. Nat. Protoc 2022, 17 (8), 1832–1867.

12. Pade, L. R.; Stepler, K. E.; Portero, E. P.; DeLaney, K.; Nemes, P., Biological mass spectrometry enables spatiotemporal ‘omics: From tissues to cells to organelles. Mass Spectrom. Rev. 2023, No. e21824.

13. Nemes, P., Mass spectrometry comes of age for subcellular organelles. Nat. Methods 2021, 18 (10), 1157–1158.

14. Behnke, J.-S.; Urner, L. H., Emergence of mass spectrometry detergents for membrane proteomics. Anal. Bioanal. Chem. 2023, 415 (18), 3897–3909.

15. Saha-Shah, A.; Esmaeili, M.; Sidoli, S.; Hwang, H.; Yang, J.; Klein, P. S.; Garcia, B. A., Single cell proteomics by data-independent acquisition to study embryonic asymmetry in *Xenopus laevis*. Anal. Chem. 2019, 91 (14), 8891–8899.

16. Martins, A. M. A.; Santos, M. D. M.; Camillo-Andrade, A. C.; Leite, A. L.; Souza, J. S.; Sánchez, S.; Muotri, A. R.; Carvalho, P. C.; Yates, J. R., III, Integrating DIA single-cell proteomics data with the DiagnoMass proteomic hub for biological insights. J. Am. Soc. Mass Spectrom. 2024, 35 (10), 2308–2314.

17. Kitata, R. B.; Yang, J. C.; Chen, Y. J., Advances in data-independent acquisition mass spectrometry towards comprehensive digital proteome landscape. Mass Spectrom. Rev. 2022, e21781.

18. Shen, B.; Pade, L.; Nemes, P., Data-independent acquisition shortens the analytical window of single-cell proteomics to fifteen minutes in capillary electrophoresis mass spectrometry. J. Proteome Res. 2025, 23, 692−703.

19. Kongpracha, P.; Wiriyasermkul, P.; Isozumi, N.; Moriyama, S.; Kanai, Y.; Nagamori, S., Simple but efficacious enrichment of integral membrane proteins and their interactions for in-depth membrane proteomics. Mol. Cell. Proteomics 2022, 21 (5), No. 100206.

20. Zhang, Z. B.; Dubiak, K. M.; Shishkova, E.; Huber, P. W.; Coon, J. J.; Dovichi, N. J., High-throughput, comprehensive single-cell proteomic analysis of *Xenopus laevis* embryos at the 50-cell stage using a microplate-based MICROFASP system. Anal. Chem. 2022, 94 (7), 3254–3259.

21. Shen, B.; Zhou, F.; Nemes, P., Real-Time Eco-AI, Electrophoresis-correlative data-dependent acquisition with AI-based data processing broadens access to single-cell mass spectrometry proteomics. Angew. Chem. Int. Ed. 2025, 64, No. e202510692.

22. Shen, B.; Pade, L. R.; Zhou, F.; Nemes, P., Subcellular mass spectrometry reveals proteome remodeling in an asymmetrically dividing (frog) embryonic stem cell. Proc. Natl. Acad. Sci. U. S. A. 2026, 123, No. e2518372123.

23. Peuchen, E. H.; Cox, O. F.; Sun, L. L.; Hebert, A. S.; Coon, J. J.; Champion, M. M.; Dovichi, N. J.; Huber, P. W., Phosphorylation dynamics dominate the regulated proteome during early *Xenopus* development. Sci. Rep. 2017, 7, No. 15647.

24. Lombard-Banek, C.; Moody, S. A.; Nemes, P., Single-cell mass spectrometry for discovery proteomics: quantifying translational cell heterogeneity in the 16-cell frog (*Xenopus*) embryo. Angew. Chem. Int. Ed. 2016, 55 (7), 2454–2458.

25. Choi, S. B.; Polter, A. M.; Nemes, P., Patch-clamp proteomics of single neurons in tissue using electrophysiology and subcellular capillary electrophoresis mass spectrometry. Anal. Chem. 2022, 94 (3), 1637–1644.

26. Shen, B.; Zhou, F.; Nemes, P., Electrophoresis-correlative ion mobility deepens single-cell proteomics in capillary electrophoresis mass spectrometry. Mol. Cell. Proteomics 2025, 24 (2), No. 100892.

27. Heasman, J., Patterning the early *Xenopus* embryo. Development 2006, 133 (7), 1205–1217.

28. Blum, M.; Ott, T., *Xenopus*: an undervalued model organism to study and model human genetic disease. Cells Tissues Organs 2018, 205 (5-6), 303–313.

29. Moody, S. A., Fates of the blastomeres of the 16-cell stage *Xenopus* embryo. Dev. Biol. 1987, 119 (2), 560–578.

30. Tajik, M.; Baharfar, M.; Donald, W. A., Single-cell mass spectrometry. Trends Biotechnol. 2022, 40 (11), 1374–1392.

31. Pade, L. R.; Lombard-Banek, C.; Li, J.; Nemes, P., Dilute to enrich for deeper proteomics: A yolk-depleted carrier for limited populations of embryonic (frog) cells. J. Proteome Res. 2024, 23 (2), 692–703.

32. Baxi, A. B.; Lombard-Banek, C.; Moody, S. A.; Nemes, P., Proteomic characterization of the neural ectoderm fated cell clones in the *Xenopus laevis* embryo by high-resolution mass spectrometry. ACS Chem. Neurosci. 2018, 9 (8), 2064–2073.

33. Jorgensen, P.; Steen, J. A. J.; Steen, H.; Kirschner, M. W., The mechanism and pattern of yolk consumption provide insight into embryonic nutrition in *Xenopus*. Development 2009, 136 (9), 1539–1548.

34. Liu, J.; Wang, F.; Mao, J.; Zhang, Z.; Liu, Z.; Huang, G.; Cheng, K.; Zou, H., High-Sensitivity N-glycoproteomic analysis of mouse brain tissue by protein extraction with a mild detergent of N-dodecyl β-D-maltoside. Anal. Chem. 2015, 87 (4), 2054–2057.

35. Sokol, S. Y., Wnt signaling and dorso-ventral axis specification in vertebrates. Curr. Opin. Genet. Dev. 1999, 9 (4), 405–410.

36. De Robertis, E. M.; Kuroda, H., Dorsal-ventral patterning and neural induction in *Xenopus* embryos. Annu. Rev. Cell Dev. Biol. 2004, 20, 285–308.

