## Supplementary material for "Interface-Resolved Proteomics of Cell–Cell Membranes Reveals Early Spatial Polarity in a Vertebrate Embryo": SI Document

### SI METHODS

**Materials and Reagents.** Liquid chromatography (LC) mass spectrometry (MS) quality solvents (acetonitrile, ACN; formic acid–FA) were purchased from Thermo Scientific or MilliporeSigma. All reagents were purchased at HPLC grade or higher: ammonium bicarbonate (AmBic), dithiothreitol (DTT), iodoacetamide (IAA), and n-dodecyl-β-D-maltoside (DDM). *Steinberg's solution (100% SS)* for embryology: 17% (w/v) sodium chloride, 0.5% (w/v) potassium chloride, 0.5% (w/v) calcium chloride, and 1.025% (w/v) magnesium sulfate. *Homogenization buffer:* 10 mM Tris–HCl, 150 mM NaCl, 250 mM sucrose, 1 mM EDTA, and protease inhibitor cocktail. *Lysis buffer*: 1% (w/v) sodium dodecyl sulfate (SDS).

**Animals.** The sexually mature *Xenopus laevis* were sourced from Nasco or Xenopus1 (Dexter, MI). The frogs were maintained and cared for humanely following protocols approved by the University of Maryland's Institutional Animal Care and Use Committee (approval nos. R-FEB-21-07 and R-FEB-24-05). The embryos were obtained from natural matings between 2 breeding pairs. The jelly coat was removed with 2% (w/v) cysteine, followed by 3 washes in 10% S.S. Embryos with stereotyped pigmentation were selected at the 2-cell stage and cultured at 14 °C in 100% SS. During a 2-h incubation, the embryos were monitored under a stereomicroscope, and those reaching the 16-cell stage were transferred to an agar-coated plate for membrane microdissection. The cells were identified based on pigmentation, size, and location in reference to established cell-fate[1] models.

**Subcellular Membrane Isolation.** The vitelline membrane was carefully torn and removed with forceps under a stereomicroscope. Two adjacent animal-hemisphere cells (D12 and V12) sharing the intercellular membrane of interest were dissected from the animal side with minimal perturbation to the target membrane. The cell-cell interfaces were then microdissected using a microscalpel and transferred into "low-analyte loss" microtube (protein LoBind, 13-698-793, Fisher Scientific) microcentrifuge tubes. For method development, membranes between D12 and V12 (DV) were collected. For biological application, the membranes from 3 locations were randomly collected from embryos derived from 4 breeding pairs across 2 natural matings to account for natural biological variability.

**Membrane Enrichment by Ultracentrifugation.** To prepare the membrane portion, the samples were manually homogenized in 250 µL homogenization buffer on ice, followed by brief sonication in an ice bath with intermittent vortexing. Cellular debris and large complexes were pelleted at 1,000 × *g* for 5 min, 4 °C. The post-1,000 × *g* supernatant was then centrifuged at 10,000 × *g* for 10 min, 4 °C to remove nuclei and mitochondria. The resulting supernatant was concentrated by vacuum centrifugation to yield a membrane fraction. For membrane enrichment, the supernatant was subjected to ultracentrifugation at 438,000 × *g* for 30 min, 4 °C using a Beckman Airfuge (see Ref.[2]). The supernatant was carefully removed, and the membrane-enriched pellet was retained for bottom-up proteomic sample preparation.

**Bottom-up Proteomic Workflow.** The dissected membrane proteome was processed for bottom-up proteomics as per the following steps. ***Lysis***. The enriched membrane pellets and dissected membrane samples were lysed in the *lysis buffer*, followed by protein precipitation with chilled acetone (≥4× volume, −20 °C, overnight). For the limited dissected membranes, this step was alternatively performed with the nonionic detergent DDM at 0.1%, 0.5%, 1%, or 2% (w/v), without precipitation. ***Reduction/Alkylation***. The extracted proteins were denatured/reduced with 20 mM DTT at 60 °C for 30 min, alkylated with 60 mM IAA for 20 min at room temperature in the dark, then quenched with 20 mM DTT. The samples were adjusted/reconstituted in 50 mM AmBic. ***Digestion***. The protein samples were digested with trypsin at 1:20 (enzyme: protein, w/w) for 12 h at 37 °C with gentle mixing. ***Fractionation.*** To reduce the sample complexity and generate the spectral library, high-PH (HPH) reversed-phase chromatography (1260 Infinity II LC systems, Agilent, Santa Clara, CA). The peptide mixtures were loaded onto a C18 guard column (2.7 µm, 4.6 × 5 mm, Agilent InfinityLab Poroshell), and separated on the HPH-C18 analytical column (2.7 µm, 4.6 × 150 mm, Agilent). HPH fractionation used 10 mM AmBic in water (pH 10) as solvent A and 10 mM AmBic in 90% ACN (pH 10) as solvent B. The pool membrane sample (~50 µg in total) were separated into 72 fractions at a flow rate of 0.5 mL/min, using a 90-min gradient of solvent B as follows, 0−17 min, 0%; 17−18 min, to 7%; 18−75 min, to 35%; 75−76 min, to 100%; 76−80 min, hold 100%; 80−86 min, to 0%; 86−90 min, hold 0% (equilibration). Fractions were vacuum-dried and concatenated into eight fractions. ***Cleanup.*** The resulting peptide mixtures were desalted on C18 spin columns (part. no. 89870, Thermo Fisher Scientific) according to the manufacturer's instructions, dried by vacuum centrifugation, and reconstituted in 0.1% (v/v) FA in LC–MS-grade water.

**nanoLC–HRMS.** Sample Preparation–Loading. ~0.5 µg of the peptides (from Total Peptide Assay, Thermo) was loaded onto a C18 trap column (100 µm inner diameter, 5 µm particle with 100 Å pores, 2 cm length, Acclaim PepMap 100, Thermo), followed by a 5 min flush at 1% buffer B at 5 µm/min for online desalting. ***Chromatography*** used 0.1% FA in water as the weak mobile phase (solvent A) and 0.1% FA in ACN as the strong (solvent B). The peptides were separated on a C18 µPAC column (200 cm, S/N 1100462, Thermo) at 600 nL/min using a 240 min gradient of B as follows: 0–5 min, 1%; 5–20 min, to 7%; 20–135 min, to 25%; 135–160 min, to 32%; 160–193 min, to 45%; 193–200 min, to 80%; 200–208 min, hold 80% (column flush); 208–210 min, ramp to 2%; and 210–240 min, 2% (equilibration).

**Ionization and MS.** The eluting peptides were ionized by electrospray (ESI) and detected on an Orbitrap Fusion Lumos Tribrid (Thermo) in the positive-ion mode. The instrument was operated under data-dependent acquisition (DDA) using the following settings: $MS^1$ scan was acquired in orbitrap with a resolution of 120,000 (Full Width at Half Maximum, FWHM at *m/z* 200); scan range, *m/z* 350–1,500; maximum injection time (max IT), 50 ms; intensity threshold, $5.0 \times 10^3$; and dynamic exclusion 60 s. Precursor selection: quadrupole isolation window, 1.6 Th. $MS^2$ scans were acquired in the ion trap with flowing settings: standard AGC target; duty cycle, 3 s total cycle time; 30% HCD collision energy. Alternatively, ***data-independent acquisition (DIA)*** $MS^1$ scans were acquired in Orbitrap with a resolution of 120,000. The $MS^2$ scans were acquired in Orbitrap with a resolution of 30,000. Schemes were evaluated with a comparable duty cycle (~3 s/cycle) using the following combinations of m/z scan range, isolation window width, and number of isolation windows, respectively: 450–900, 10×45; 400–1,100, 15×47; 350–1,500, 25×46.

**Data Analysis.** ***DDA*** (Proteome Discoverer, PD 3.0). Database: species, *Xenopus laevis;* source, UniProt; FASTA content, 43,236 entries; date, downloaded May 4, 2022). Enzyme digestion: specificity, trypsin; max missed cleavages, 2; minimum peptide length, 5 amino acids; Modifications: carbamidomethyl (C), static; oxidation (M), variable (max variable modifications per peptide, 1). Mass tolerances: precursor, 10 ppm; fragment, 0.06 Da. FDR control, <1% using a target-decoy strategy (reversed sequence proteome). Other settings: PD defaults unless specified above. Common human contaminants were excluded by the common Human Contaminant FASTA database (cRAP−Contaminants of Resource for Affinity Purification).[3]

**DIA** (DIA-NN v19.0[4]). Search mode: DIA data processed in library-based mode using the spectral library described below; library-free analyses were also performed where indicated. Enzyme: specificity, trypsin; max missed cleavages, 2. Modifications: carbamidomethyl (C), static; oxidation (M), variable (max variable modifications per peptide, 1). Peptide length range: 5–25 amino acids. Precursor charge range: +2–4. FDR control: <1% (DIA-NN defaults for q-value estimation). Other settings: DIA-NN defaults unless specified above. **Protein–protein associations** were queried in STRING 12.0[5] with the following settings: disconnected nodes, hidden; clustering, k-means; number of clusters, 5.

**Data Availability.** The primary and processed MS files, MS spectral library, and *X. laevis* reference proteome were deposited in the ProteomeXchange Consortium via the PRIDE partner repository under accession number identifier PXD067644 (https://www.ebi.ac.uk/pride/).

**Safety.** All procedures were performed in accordance with institutional chemical hygiene plans and animal care protocols. Personnel received appropriate training and consulted Safety Data Sheets for all reagents and solvents.

**Scientific rigor**. For spatial analyses, intercellular membranes from the DD, DV, and VV interfaces at the 16-cell stage were collected as 4–5 biological replicates; for each region, n = 3 membranes were pooled per biological replicate from embryos derived from 2 independent natural matings. Unless noted otherwise, each pooled digest was analyzed by LC–MS in technical triplicate. Relative protein abundance was estimated from label-free quantification (LFQ) intensities. LFQ values were sum-normalized and $\log_{10}$-transformed, before multivariate analyses; principal component analysis and hierarchical cluster analysis were performed on the autoscaled data. Differential abundance between regions was tested with 2-sided Student's t-tests, with $p < 0.05$ considered significant from MetaboAnalyst 6.0.[6] Reported $p$-values are nominal (no multiple-testing correction) unless otherwise specified in the figure or legend. Select illustrations were created with help from BioRender.com.

### SI REFERENCES

1. Moody, S. A., Fates of the blastomeres of the 16-cell stage *Xenopus* embryo. *Dev. Biol.* **1987,** *119* (2), 560-578.
2. Kongpracha, P.; Wiriyasermkul, P.; Isozumi, N.; Moriyama, S.; Kanai, Y.; Nagamori, S., Simple but efficacious enrichment of integral membrane proteins and their interactions for in-depth membrane proteomics. *Molecular & Cellular Proteomics* **2022,** *21* (5), No. 100206.
3. Mellacheruvu, D.; Wright, Z.; Couzens, A. L.; Lambert, J. P.; St-Denis, N. A.; Li, T.; Miteva, Y. V.; Hauri, S.; Sardiu, M. E.; Low, T. Y.; Halim, V. A.; Bagshaw, R. D.; Hubner, N. C.; Al-Hakim, A.; Bouchard, A.; Faubert, D.; Fermin, D.; Dunham, W. H.; Goudreault, M.; Lin, Z. Y.; Badillo, B. G.; Pawson, T.; Durocher, D.; Coulombe, B.; Aebersold, R.; Superti-Furga, G.; Colinge, J.; Heck, A. J. R.; Choi, H.; Gstaiger, M.; Mohammed, S.; Cristea, I. M.; Bennett, K. L.; Washburn, M. P.; Raught, B.; Ewing, R. M.; Gingras, A. C.; Nesvizhskii, A. I., The CRAPome: a contaminant repository for affinity purification-mass spectrometry data. *Nat. Methods* **2013,** *10* (8), 730-736.

4. Demichev, V.; Messner, C. B.; Vernardis, S. I.; Lilley, K. S.; Ralser, M., DIA-NN: neural networks and interference correction enable deep proteome coverage in high throughput. *Nat. Methods* **2020,** *17* (1), 41-44.
5. Szklarczyk, D.; Nastou, K.; Koutrouli, M.; Kirsch, R.; Mehryary, F.; Hachilif, R.; Hu, D.; Peluso, M. E.; Huang, Q.; Fang, T.; Doncheva, N. T.; Pyysalo, S.; Bork, P.; Jensen, L. J.; von Mering, C., The STRING database in 2025: protein networks with directionality of regulation. *Nucleic Acids Res.* **2025,** *53* (D1), D730-D737.
6. Pang, Z.; Lu, Y.; Zhou, G.; Hui, F.; Xu, L.; Viau, C.; Spigelman, Aliya F.; MacDonald, Patrick E.; Wishart, David S.; Li, S.; Xia, J., MetaboAnalyst 6.0: towards a unified platform for metabolomics data processing, analysis and interpretation. *Nucleic Acids Research* **2024,** *52* (1), 398-406.

### SI FIGURES

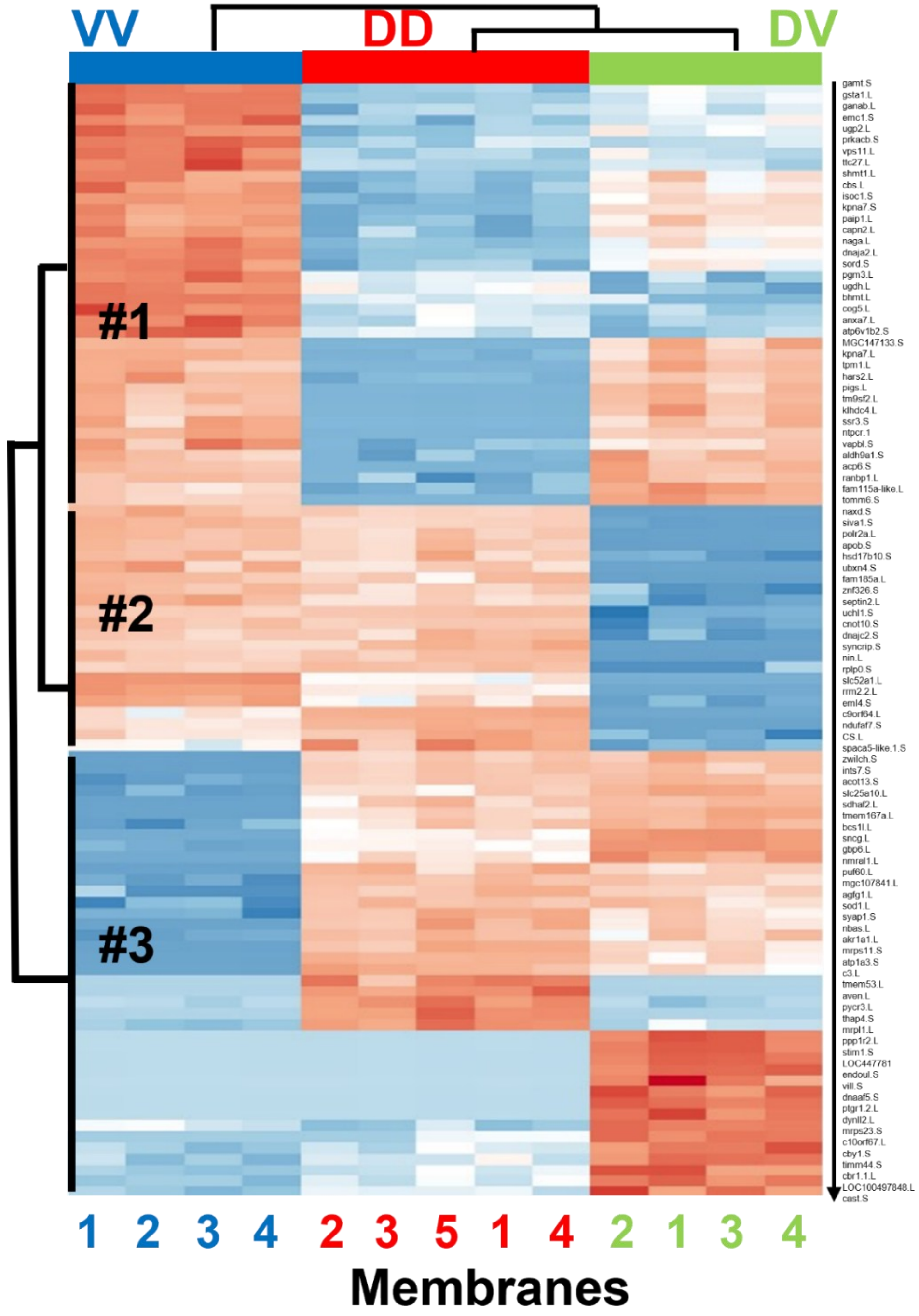

**Figure S1.** Heat map of the top 100 varying total proteins across regions (rows, z-scored LFQ; columns, biological replicates). Color code to heat maps (rows, z-scored LFQ): higher abundance (red), lower abundance (blue).

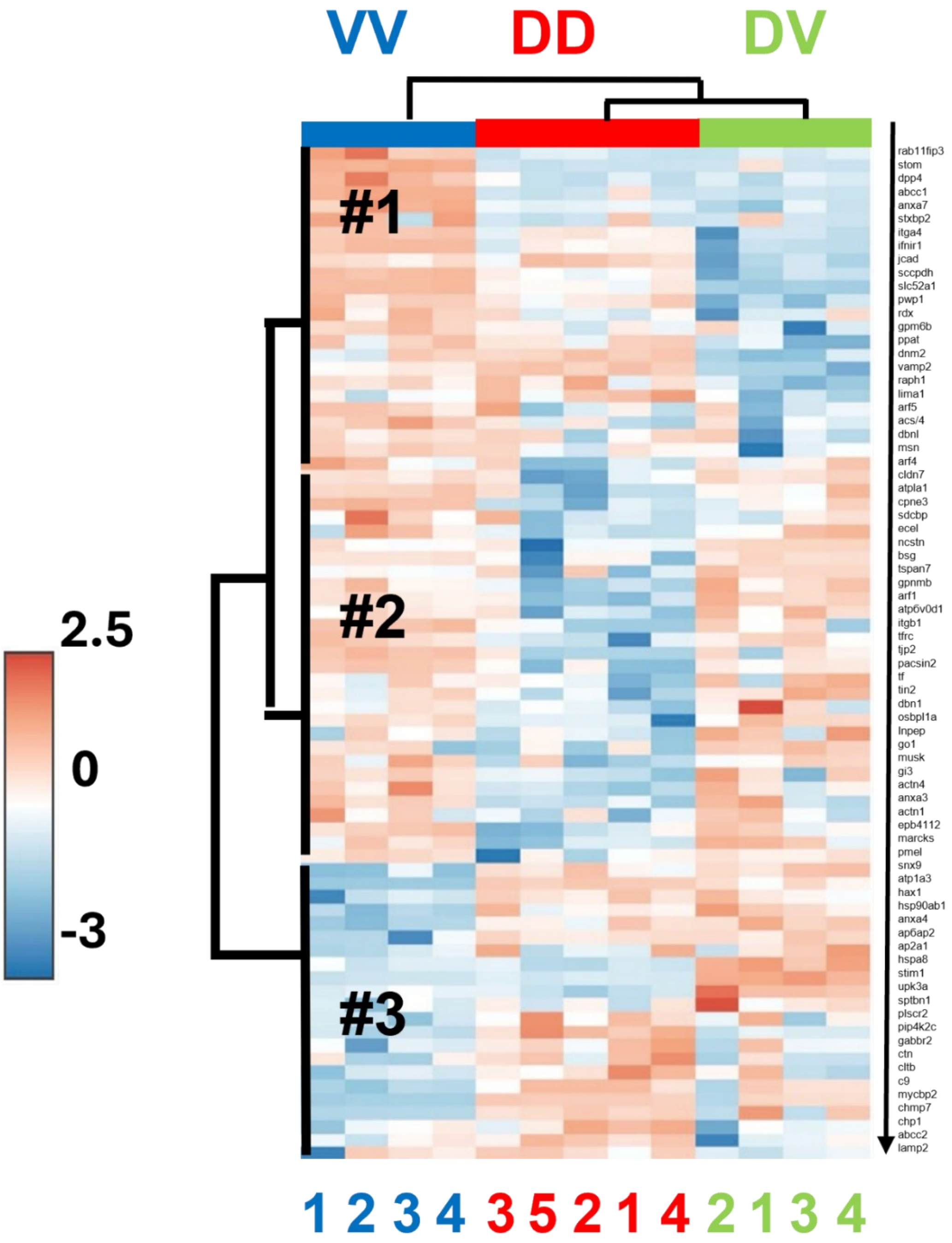

**Figure S2.** Heat map of the top 75 plasma-membrane proteins (rows) across DD, DV, and VV membrane samples (columns). Color code to heat maps (rows, z-scored LFQ): higher abundance (red), lower abundance (blue).
